# Genetic editing of SEC61, SEC62, and SEC63 abrogates human cytomegalovirus US2 expression in a signal peptide-dependent manner

**DOI:** 10.1101/653857

**Authors:** Anouk B.C. Schuren, Ingrid G.J. Boer, Ellen Bouma, Robert Jan Lebbink, Emmanuel J.H.J. Wiertz

**Affiliations:** Department of Medical Microbiology, University Medical Center Utrecht, 3584CX Utrecht, The Netherlands; Department of Medical Microbiology, University Medical Center Groningen, Postbus 30001, 9700 RB Groningen, The Netherlands

**Author notes:** Correspondence (E.J.H.J.W.).

## Abstract

Newly translated proteins enter the ER through the SEC61 complex, via either co- or post-translational translocation. In mammalian cells, few substrates of post-translational SEC62- and SEC63-dependent translocation have been described. Here, we targeted all components of the SEC61/62/63 complex by CRISPR/Cas9, creating knock-outs or mutants of the individual subunits of the complex. We show that functionality of the human cytomegalovirus protein US2, which is an unusual translocation substrate with a low-hydrophobicity signal peptide, is dependent on expression of not only SEC61α, -β, and -γ, but also SEC62 and SEC63, suggesting that US2 may be a substrate for post-translational translocation. This phenotype is specific to the US2 signal peptide.

## Introduction

Up to one-third of all proteins is secreted or expressed in cellular or organelle membranes(Chen, Karnovsky, Sans, Andrews, & Williams, 2010; Wallin & Heijne, 2008). These proteins have been translocated into the endoplasmic reticulum (ER) during or after their translation. Translocation of newly translated proteins into the ER is facilitated by the SEC61 complex, which consists of a multimembrane-spanning SEC61α-subunit associated with smaller SEC61β and -γ subunits(Kalies, Stokes, & Hartmann, 2008). Depending on the mode of translocation, the complex can be complemented by SEC62 and SEC63(Deshaies, Sanders, Feldheim, & Schekman, 1991; A. E. Johnson & van Waes, 1999; Linxweiler, Schick, & Zimmermann, 2017).

Translocation of proteins into the ER can occur either co- or post-translationally(Rapoport, 2007; Zimmermann, Eyrisch, Ahmad, & Helms, 2011). The canonical co-translational route is well established in both yeast and mammalian cells. During translation of a signal peptide, the signal recognition particle (SRP) binds the translating ribosome and guides it towards the SRP receptor at the ER membrane. The SRP receptor subsequently interacts with SEC61α, such that the nascent chain is translocated over the ER membrane as translation continues(Song, Raden, Mandon, & Gilmore, 2000). Lateral opening of the SEC61 complex allows for release of translated transmembrane domains(A. E. Johnson & van Waes, 1999).

Post-translational translocation remains elusive in higher eukaryotes but is well-studied in yeast. Cytosolic recognition of post-translationally translocated proteins occurs independently of SRP. Instead, cytosolic chaperones of the Hsp70- and Hsp40 families guide fully translated proteins towards the ER(Chirico, Waters, & Blobel, 1988). Signal sequence characteristics determine which mode of translocation is used. Although signal peptides do not share sequence identity, there is overall homology with regard to their polarity and hydrophobicity(von Heijne, 1985). All signal sequences contain an amino-terminal n-region, a central h-region, and a carboxyterminal c-region, containing the cleavage site for removal of the signal peptide. The n- and c-regions contain charged and polar residues, whereas the h-region is hydrophobic(von Heijne, 1985). Post-translational translocation occurs mostly for proteins with modestly hydrophobic signal sequences because they interact less strongly with SRP(Karamyshev et al., 2014; Ng, Brown, & Walter, 1996; Nilsson et al., 2015; Reithinger, Kim, & Kim, 2013; Zheng & Nicchitta, 1999), and for signal sequences with a positive charge in the n-region(Guo et al., 2018). Post-translational translocation has been suggested to act as a back-up mechanism for ER delivery of proteins that fail to translocate via SRP, either because of suboptimal signal sequence functioning(Conti, Devaraneni, Yang, David, & Skach, 2015; Guo et al., 2018; Kriegler, Magoulopoulou, Amate Marchal, & Hessa, 2018), or because of their short length. Proteins shorter than 110 amino acids in length are translocated in a post-translational manner. It is speculated that translation of these short proteins occurs too rapidly to allow for co-translational recognition by SRP, suggesting that they are recognized post-translationally(N. Johnson, Powis, & High, 2013; Lakkaraju et al., 2012).

Another feature that influences translocation efficiency is the usage frequency of amino acid codons in the signal peptide. Signal peptides in general contain a high number of sub-optimal codons. A low concentration of tRNAs binding these rare codons slows down translation, allowing more time for the nascent chain to engage SRP(Pechmann, Chartron, & Frydman, 2014; Wang, Mao, Xu, Tao, & Chen, 2015; Yu et al., 2016; Zalucki, Beacham, & Jennings, 2009), thereby improving (co-translational) translocation efficiency(Guo et al., 2018). Correspondingly, slowing down translation using a low dose of cycloheximide strongly increases translocation efficiency(Guo et al., 2018), while replacing all sub-optimal codons in a signal sequence by their optimal counterparts, thereby accelerating translation, can lower protein expression at least 20-fold(Zalucki, Jones, Ng, Schulz, & Jennings, 2010).

In yeast, post-translational translocation is mediated by an auxiliary complex, containing the Sec61α/β/γ complex extended with Sec62p, Sec63p, Sec71p and Sec72p(Brodsky & Schekman, 1993; Deshaies et al., 1991; Zimmermann et al., 2011). The structure of this complex has recently been solved(Itskanov & Park, 2019; Wu, Cabanos, & Rapoport, 2019). In mammalian cells, the homologous SEC62 and SEC63 have been suggested to play a role in post-translational translocation(Lang et al., 2012). SEC63 is a Hsp40 chaperone which, via its J-domain, interacts with the ER chaperone BiP to provide the driving force for translocation(Lyman & Schekman, 1995). SEC62 provides an alternative for SRP-mediated translocation. The SEC61 complex can switch from SRP-dependent to SEC62-dependent translocation, when SEC62 outcompetes the SRP receptor for SEC63 binding(Jadhav et al., 2015).

The human cytomegalovirus (HCMV) protein US2 is implicated in evading immune recognition via degradation of immunoreceptors including HLA class I via ER-associated protein degradation (ERAD) (Hsu et al., 2015; Wiertz, Tortorella, et al., 1996). US2 contains an atypical signal peptide with low hydrophobicity(Gewurz, Ploegh, & Tortorella, 2002), resulting in inefficient translocation that is potentially facilitated by SEC62 and SEC63(Conti et al., 2015). Here, we assessed which mode of translocation is co-opted by US2. We established clonal cell lines with in-frame or out-of-frame indels for each SEC61/62/63 component. We report that SEC61, SEC62 and SEC63 components are essential for translocation of US2, using US2-mediated ERAD of HLA-I as a readout. The weak signal sequence may direct US2 to the SEC62/SEC63 post-translational translocation pathway. This effect is specific to the US2 signal peptide and the signal peptide is sufficient to sensitize a protein to SEC61/62/63 editing. The SEC mutants obtained provide insights into the structural and functional roles of several of these proteins.

## Results

### CRISPR/Cas9-mediated editing of all SEC61/62/63 subunits compromises US2 translocation

The HCMV-encoded immune evasion protein US2 is an unconventional translocation substrate that has a signal sequence with low hydrophobicity. CRISPR/Cas9 technology enabled us to study US2 in the context of genetically mutated SEC61 subunits. US2-mediated HLA class I degradation was used as an indirect readout for US2 translocation. CRISPR single guide RNAs (sgRNAs) targeting subunits of the SEC61 complex were introduced in a clonal U937 cell line expressing HLA-A2-eGFP, US2 and Cas9. As a control, we CRISPR-targeted the ubiquitin E3 ligase TRC8, which is known to be essential for US2-mediated HLA-I downregulation(Stagg et al., 2009).

Flow cytometry was used to determine the expression levels of HLA-A2 as a measure of US2 function. As it is known that ER-resident HLA-I that is rescued from degradation can either accumulate in the cytosol or escape to the cell surface(Stagg et al., 2009; van de Weijer et al., 2017), we measured both the total HLA-A2 expression by means of its eGFP-tag, as well as cell surface expression using an HLA-A2-specific antibody (fig. 1). We tested sgRNAs targeting all subunits of the SEC61/62/32 complex and observed HLA-A2 rescue for SEC61α1, SEC61β, SEC61γ, SEC62 and SEC63 (fig. 1), indicating that US2 function is disrupted in these cells. These data are in agreement with a genome-wide CRISPR/Cas9 library screen we have performed on HLA-I rescue in the presence of US2 (manuscript in preparation). Only SEC61α2 knock-out did not hamper US2 function. Few biochemical studies have been performed on SEC61α2 to date, and it is unclear whether this subunit is in fact part of a functional translocation complex in U937 cells.

**Figure 1:**
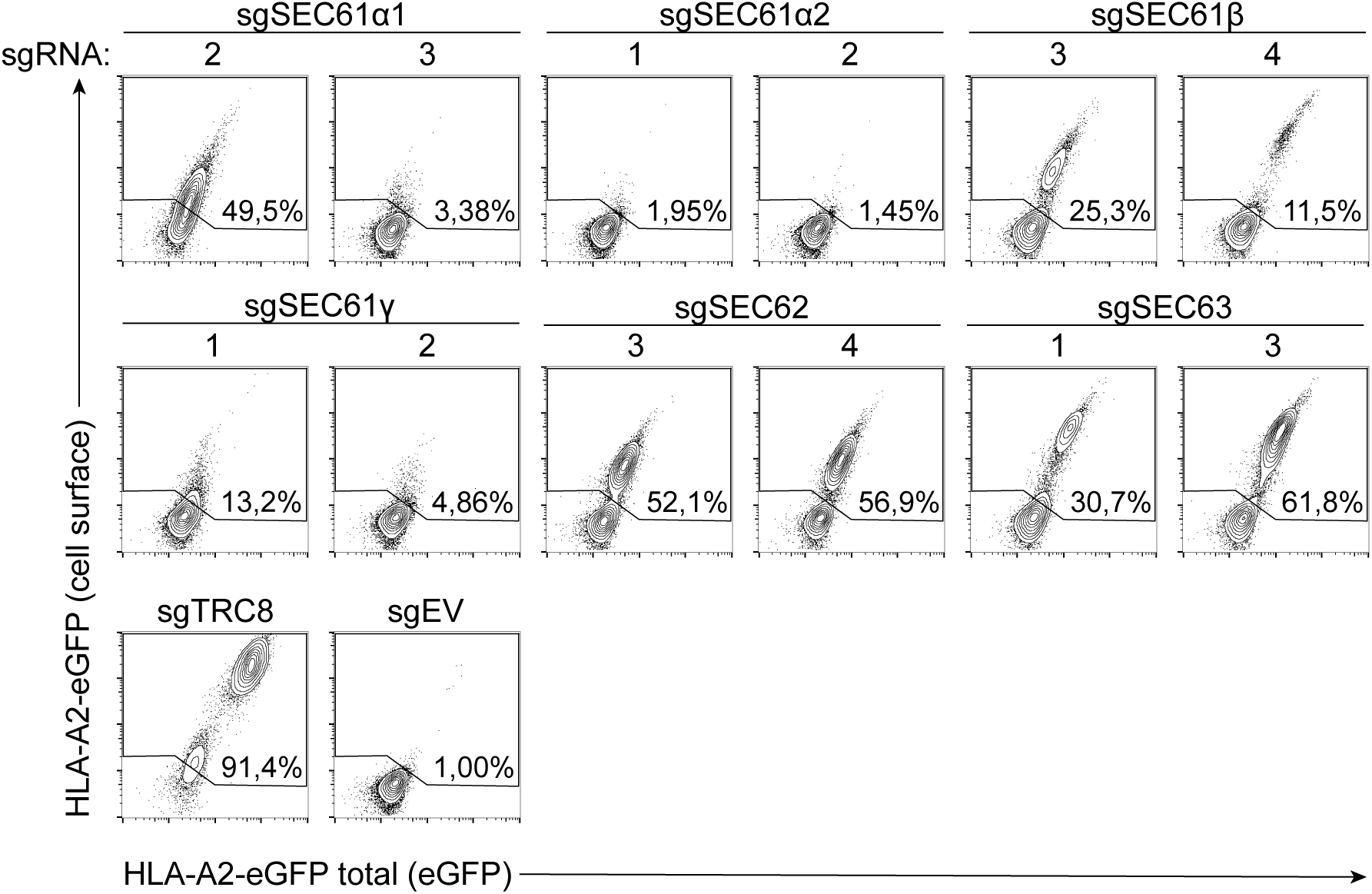
The SEC61 complex plays a role in US2-mediated degradation of HLA-I. CRISPR/Cas9 sgRNAs targeting SEC61 subunits, or control sgRNAs, were added to U937 cells expressing HLA-A2-eGFP and untagged US2. Four sgRNAs were introduced per gene, of which the two most potent are shown. At 12 days post-infection, HLA-A2 expression was assessed by flow cytometry. Total (intracellular and extracellular) HLA-A2 expression was measured by means of its eGFP-tag. Cell surface expression of HLA-A2-eGFP was assessed by an HLA-A2-specific (BB7.2) antibody staining. sgRNA sequences are listed in the materials and methods. All knock-outs of SEC61 subunits, with the exception of SEC61α2, rescue HLA-I from US2-mediated degradation.

The clear HLA-A2 rescue phenotype at 12 days post-transduction with the SEC sgRNAs (fig. 1) was surprising. Our experience is that knocking out essential ERAD genes, such as p97/VCP and its co-factors Ufd1 and Npl4, yields a minor and transient HLA-I rescue phenotype due to lethality of the knock-out cells and hence loss of the phenotype over time (fig. 2A). By contrast, measuring the SEC61-edited cells over time showed that the HLA-A2 rescue phenotypes were apparent for at least 28 days (fig. 2B), although the effect waned after a peak at 11-14 days post-transduction. The phenotype also seemed specific to the HCMV protein US2, as the activity of another ERAD-inducing HCMV protein, US11(Wiertz, Jones, et al., 1996), was not sensitive to SEC61 editing (fig. 2C), with the exception of SEC63.

**Figure 2:**
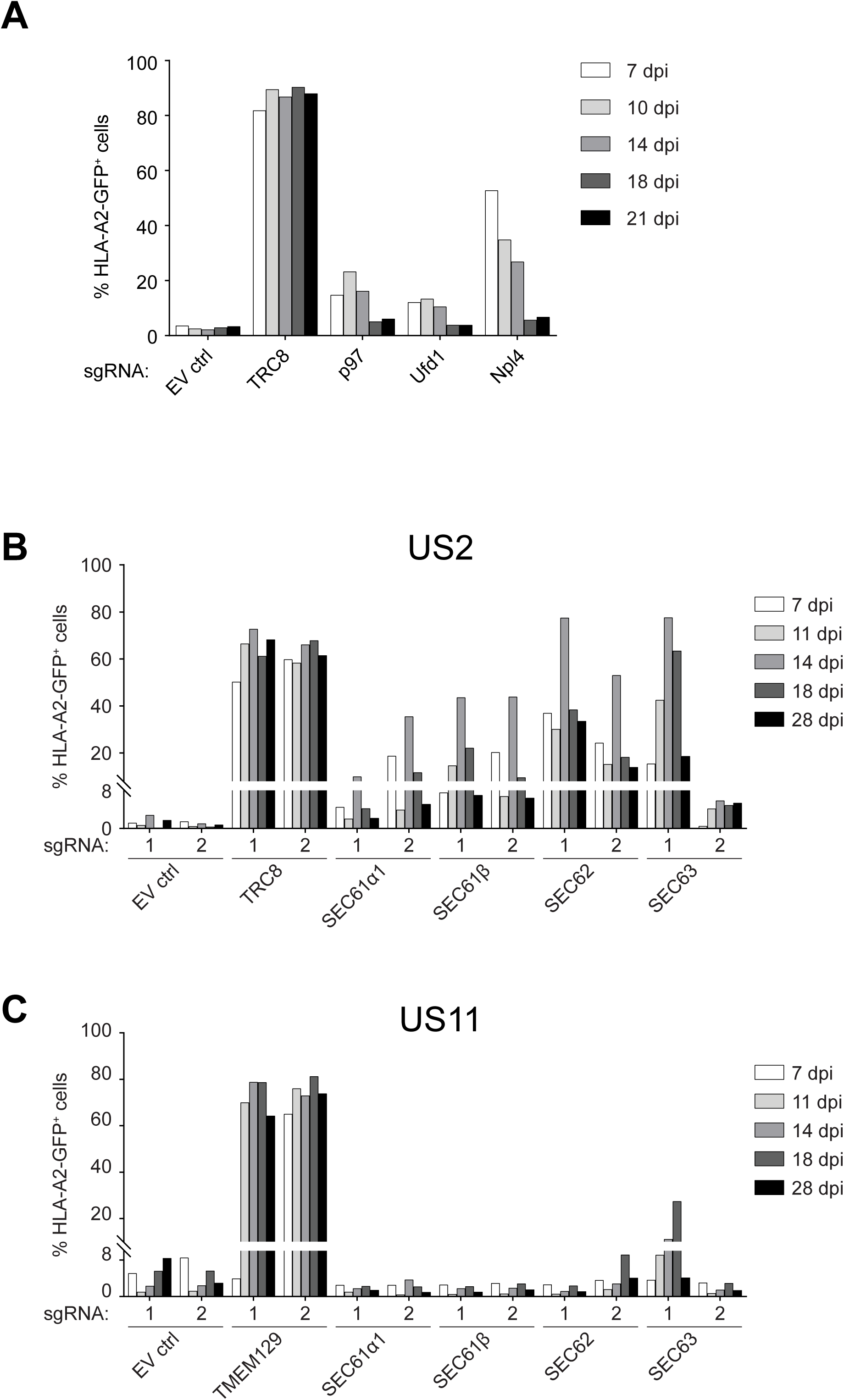
CRISPR/Cas9 targeting of SEC61 subunits yields stable abrogation of US2-mediated HLA-I degradation. (A) Addition of CRISPR/Cas9 sgRNAs targeting essential genes such as p97/VCP or its co-factors Ufd1 and Npl4 leads to a minor and temporary rescue of HLA-I expression which is lost over time due to lethality of the knock-out. sgRNAs targeting p97, Ufd1, Npl4, TRC8 (positive control), or an empty vector lacking the sgRNA were added to HLA-A2-eGFP- and US2-expressing U937 cells. Expression of HLA-A2-eGFP was assessed in flow cytometry by means of its eGFP tag at the timepoints indicated. (B) Using a similar set-up as in A, sgRNAs targeting SEC61α1, SEC61β, SEC62, SEC63, TRC8 (positive control) or the empty vector control were added to HLA-A2-eGFP- and US2-expressing U937 cells. Targeting these SEC61 subunits yields a stable HLA-I rescue phenotype over at least 28 days. This phenotype is specific to the HCMV protein US2, since US11 (C), another ERAD-inducing HCMV protein is not sensitive to SEC61 knock-out. C) Similar to B, except that U937 cells expressing US11 instead of US2 were used.

### Stable mutant cell lines can be generated for all SEC61 components

Since the HLA-I rescue phenotype in SEC61 knock-out cells declined at later timepoints (fig. 2B) we next assessed whether this was caused by cell death due to SEC61 subunit knockout, or by a growth disadvantage of the mutant cells over non-targeted cells in the polyclonal cell population. In the latter case, clonal cell lines with a SEC61 mutation might survive long-term and enable us to study their effect on US2 functioning. For this we sorted single HLA-A2^high^ cells from the polyclonal SEC61/62/63 mutant populations by FACS and allowed these to grow out. We were able to establish clonal lines that displayed stable rescue of HLA-I (fig. 3A). CRISPR/Cas9-induced mutations in these clonal cell lines were assessed by sequencing the sgRNA target regions (table S1). The majority of the clones showed in-frame indels in one or both alleles (table S1 and fig. S1), although full genetic knock-outs of SEC61β, SEC61γ and SEC62 were selected with out-of-frame deletions in both alleles. For each SEC61/62/63 subunit, two clones were selected for further characterization (marked with ‘1’ and ‘2’ in fig. S2). All clones showed stable rescue of HLA-A2 (fig. 3A), although some displayed a decline in growth rate (data not shown).

**Figure 3:**
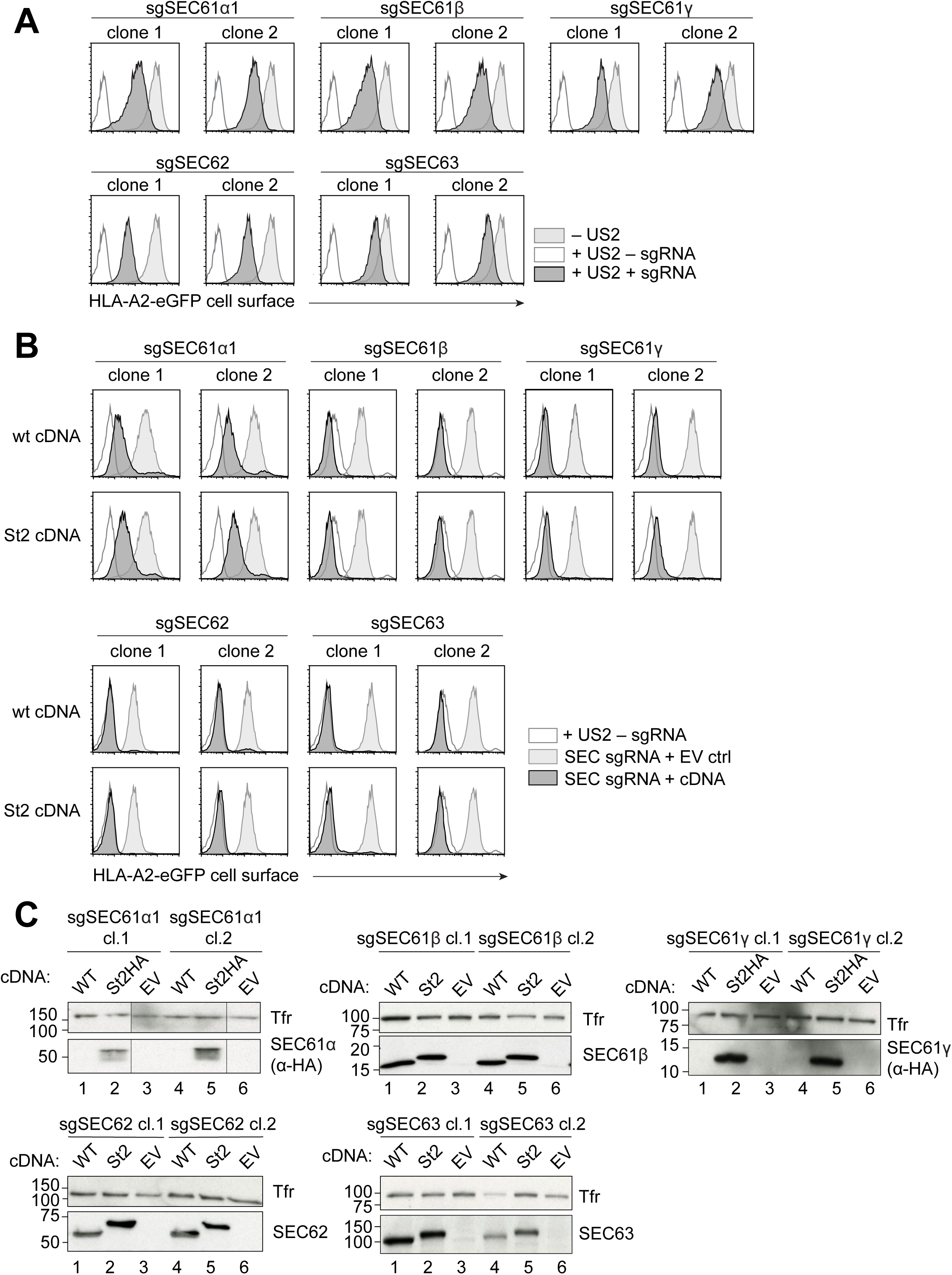
SEC61/62/63 mutant clonal lines are stable and interfere with US2-mediated HLA-I expression. A) Single cells from fig. 2B were isolated by FACS and two clonal lines were selected per edited gene, based on viability, HLA-I rescue potency (see also figure S1) and sequencing of the sgRNA target sites (table S1). HLA-A2 expression was assessed in these clonal cell lines and compared to parental cell lines without a SEC sgRNA or without US2. Selected clones yield a stable HLA-I rescue phenotype (compare black peaks to non-filled gray peaks). B) sgRNA-resistant wildtype or StrepII-(HA-)tagged cDNAs were transduced in the respective mutant clones and were selected for using Zeocin treatment. Addition of a sgRNA-resistant cDNA reverts the mutant phenotypes, showing that the HLA-I rescue effects are specific. C) Lysates of all selected mutant clones shown in A were immunoblotted and stained against the sgRNA-targeted gene. Transferrin receptor (TfR) was added as a loading control.

### SEC61/62/63 cDNAs revert the HLA-I rescue phenotype of mutant clones

To ascertain that the HLA-I rescue phenotypes we observed were specific to the SEC61/62/63 genes that were edited, we expressed sgRNA-resistant cDNAs of the respective genes in the clonal mutant lines and assessed HLA-I levels by flow cytometry. HLA-I rescue phenotypes as well as growth defects were reverted by supplementing with either wildtype or StrepII-tagged cDNAs of the genes that were knocked out (fig. 3B). Expression of the cDNAs was confirmed in Western blot (fig. 3C). In clonal cell lines with a SEC61β, SEC62 or SEC63 mutation, the respective proteins were not detectable by Western blot, suggesting that gene-editing resulted in a full knockout of protein expression. The respective cDNAs could be readily detected when these were introduced. We did not observe a change in SEC61α expression in the mutant clones. To be able to detect the expression of the cDNA that was added, we tagged this cDNA with an additional HA-tag. The lack of phenotype on the SEC61α protein level may be due to the in-frame mutations we identified in the two isolated clones (table S1). Besides the two in-frame SEC61α1-edited genes, we did isolate out-of-frame clones, but these cells were unstable and died within a few weeks (data not shown). This suggests that SEC61α1 is essential for cell survival, which is in agreement with RNAi studies reported previously(Lang et al., 2012). We were unsuccessful to reliably detect SEC61γ by Western blot. Therefore, a StrepII-HA-tagged cDNA was used to detect the SEC61γ cDNA via its HA-tag, similar to the approach used for SEC61α1. Expression of all cDNAs was confirmed in Western blot.

### Genetic editing specifically affects protein expression of the targeted SEC61/62/63 component

We also tested whether the stability of non-targeted subunits was destabilized by CRISPR/Cas9-mediated editing of single subunits, as we hypothesized that the incomplete and potentially aberrantly assembled SEC61/62/63 may be recognized by protein quality-control mechanisms. As we did not assess the interactions between the SEC61/62/63 subunits, we cannot distinguish between disintegration of the SEC61/62/63 complex and the stabilization of independent subunits. We do however observe that protein expression of non-targeted subunits remains intact, suggesting that the remaining subunits of the complex are not degraded (fig. 4A).

**Figure 4:**
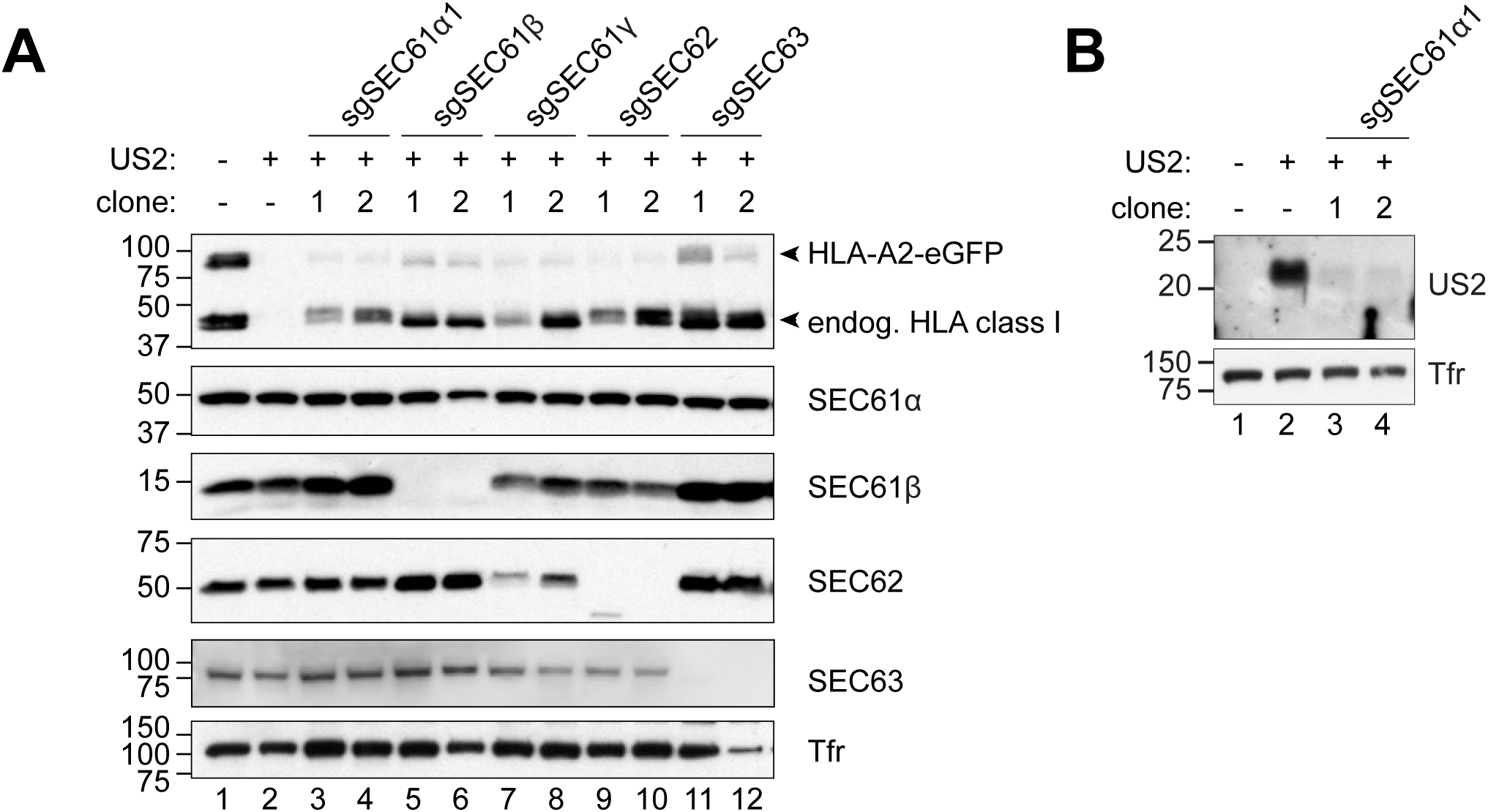
Mutating SEC61/62/63 components lowers expression of mature US2. A) Lysates of all selected mutant clones shown in fig. 3 were immunoblotted and stained against HLA class I and all components of the SEC61/62/63 complex. Transferrin receptor (TfR) was added as a loading control. Addition of CRISPR/Cas9 sgRNAs only affects expression of the targeted SEC61 subunits, but does not impact the stability of the of other subunits, suggesting that editing single subunits does not result in an unstable SEC61/62/63 complex. B) The cell lines shown in figure 3 were lysed in Triton X-100 in an experiment separate from the one shown in A. Lysates were stained with an anti-US2 antibody and an anti-transferrin receptor antibody as a loading control. A decline in US2 expression is observed upon sgRNA-targeting of SEC61α1. Preliminary data shows similar results for the other SEC subunits. Due to low US2 baseline expression, this observation could however not be repeated for these other cell lines.

In agreement with flow cytometric analysis (fig. 3A), all isolated clones expressed increased levels of HLA-A2-eGFP compared to US2-expressing control cells (fig. 4A, top panel). Endogenous HLA-I rescue was more pronounced than that of the chimeric HLA-A2-eGFP molecule (fig. 4A, bottom part of top panel).

### Expression of US2 decreases upon genetic editing of SEC61α1

We observed a strong downregulation of full-length US2 in clonal SEC61α1 mutant cells (fig. 4B), which suggests that the HLA-I rescue phenotypes are caused by a defect in US2 expression, rather than impaired ERAD functioning. Preliminary data suggest that a similar reduction in expression of mature US2 can be observed for the other clones (data not shown). The low baseline expression level of US2, even in control cells that were not targeted by CRISPR/Cas9, posed a technical difficulty that did not allow for validating the US2 expression in the other mutant clones. Expression of the transferrin receptor, another transmembrane protein that is translocated by the SEC61 complex, is not affected in our mutant clones, suggesting that this translocation defect is not a general phenomenon.

### The signal peptide determines susceptibility of US2 to SEC61/62/63 editing

To be able to detect US2 more effectively, we designed a panel of HA-tagged US2 constructs with engineered signal sequences (figs. 5A, S3 and S4). As a non-glycosylated US2 has been described(Gewurz et al., 2002; Wiertz, Tortorella, et al., 1996), potentially arising from inefficient translocation, and as the signal sequence of a protein influences its translocation efficacy(Kim, Mitra, Salerno, & Hegde, 2002), we tested the US11- and CD8 leader peptides (US11L and CD8L) alongside the original US2L.

**Figure 5:**
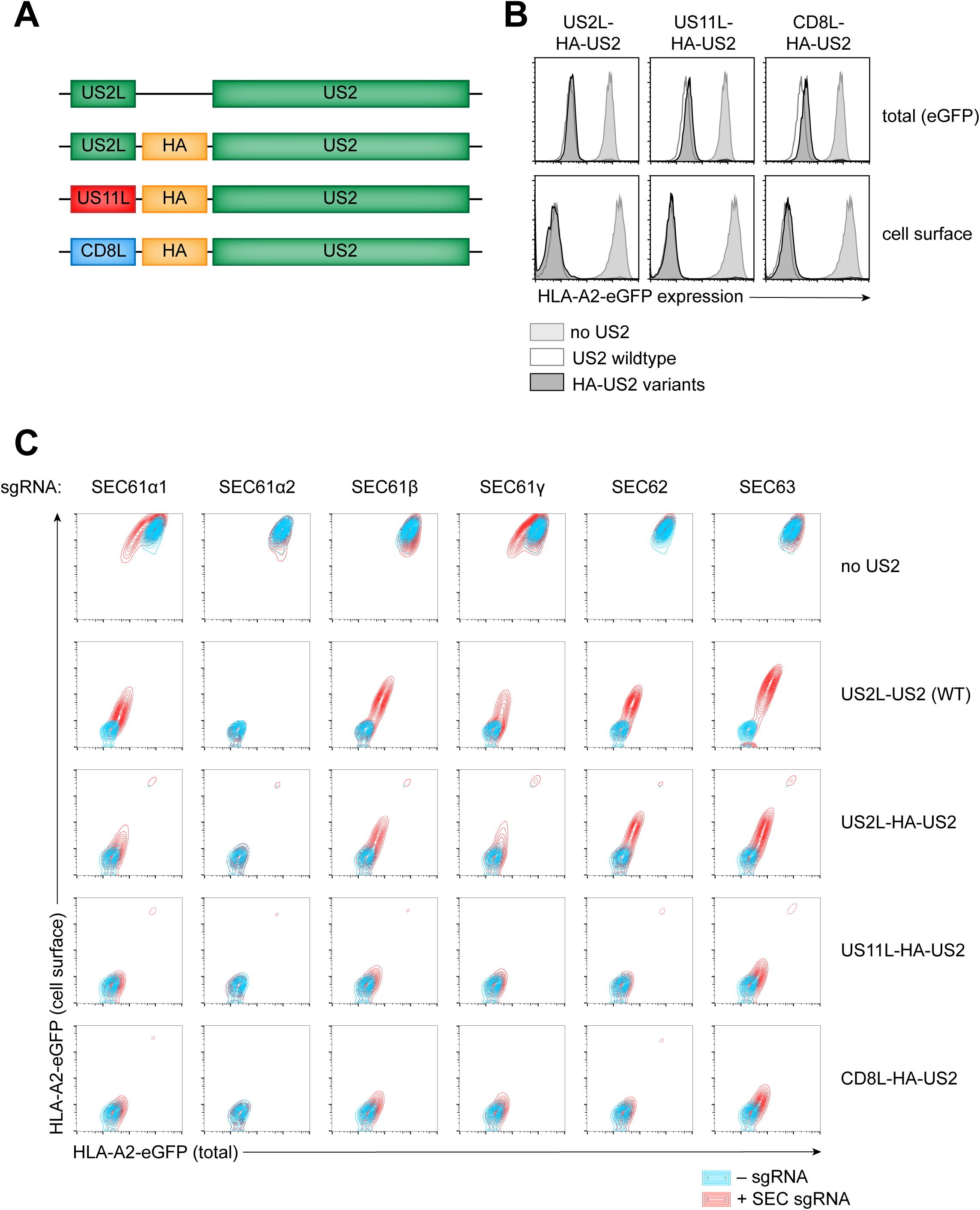
The signal peptide determines susceptibility of US2 to SEC61/62/63 editing. A) Schematic overview of HA-tagged US2-variants with different signal peptides, as compared to wildtype US2 (top). B) U937 cells expressing HLA-A2-eGFP were transduced with the US2 constructs shown in A. HLA-A2 expression was measured in flow cytometry, by means of either its eGFP-tag and an HLA-A2-specific antibody staining using BB7.2. All HA-tagged US2 variants downregulate HLA-A2 efficiently. C)U937 cells expressing HLA-A2-eGFP were transduced with the indicated HA-US2 construct or the wildtype (untagged) US2. Subsequently, CRISPR sgRNAs targeting the subunits of the SEC61/62/63 complex were introduced. This polyclonal cell population was subjected to flow cytometry to assess total levels (eGFP, x-axis) and cell-surface levels (antibody stain, Y-axis) of the HLA-A2-eGFP chimera. sgRNAs that were used in this figure are: SEC61α1 sgRNA 2; SEC61α2 sgRNA 1; SEC61β sgRNA 2; SEC61γ sgRNA 1; SEC62 sgRNA 2; SEC63 sgRNA 1.

First, we tested whether the different US2 variants were able to downregulate the HLA-A2-eGFP construct. All US2 variants were equally able to strongly downregulate HLA-A2-eGFP, as was assessed by flow cytometry (fig. 5B). We next transduced these US2-expressing cells with SEC61/62/63-targeted CRISPR sgRNAs (fig. 5C). Overlays of the flow cytometry data from cells with (red) and without (blue) sgRNAs show that only US2L-expressing US2 variants show significant HLA-A2-eGFP rescue upon SEC61/62/63 editing, whereas US2 with a CD8 or US11 leader sequence did not. This phenotype was most pronounced when targeting SEC61α1, SEC62 and SEC63. In control cells lacking US2, a minor reduction of HLA-I expression is observed only upon sgRNA-targeting of SEC61α1 and SEC61γ (fig. 5C, top panels), suggesting that HLA class I expression is also slightly hampered in these cells.

### US11 requires functional SEC61/62/63 upon transfer of the US2 signal peptide

We also tested these different signal peptides fused to the HA-tagged HCMV protein US11, which was not sensitive to SEC61/62/63 editing (fig. 2C). To be able to compare the panel of US2 variants to US11, we stably expressed a non-tagged HLA-A2 molecule in U937 cells using lentiviral transduction, as the C-terminal eGFP-tag on the HLA-A2-eGFP construct renders it insensitive to US11-mediated degradation(Cho, Kim, Ahn, & Jun, 2013). We subsequently introduced the various HA-tagged US2 and US11 variants and assessed their potential to downregulate HLA-A2 by flow cytometry (fig. 6A). All US2- and US11 variants were able to downregulate HLA-A2, with US11L- and CD8L-HA-US11 being the most potent (fig. 6A). We observed strong signal peptide-specific differences in the expression levels of the US2- and US11 variants (fig. 6B), which show an inverse correlation with the magnitude of HLA-I downregulation. Subsequently, sgRNAs targeting the SEC61/62/63 subunits were introduced in the six US2- or US11-expressing cell lines, and HLA-A2 expression was assessed by flow cytometry. To be able to compare between cell lines, HLA-A2 rescue was normalized to that in sgRNA-targeted TRC8 (for US2) or TMEM129 (for US11) cells, which were used as positive controls. The US2 variants showed an HLA-A2 rescue pattern that was similar to that observed for the HLA-A2-eGFP construct (fig. 5C), with the US2L-variant showing a substantial HLA-A2 rescue effect upon SEC61/62/63 editing, and the US11L- and CD8L-variants being less strongly affected (fig. 6C, top part). Both US11L-HA-US11 and CD8L-HA-US11 retained their ability to downregulate HLA-A2 in the presence of SEC61/62/63 sgRNAs (fig. 6C, bottom part), as was expected from studies performed in fig. 2C. Interestingly, when the US11 signal peptide was replaced by that of US2, the US11 protein gained sensitivity to SEC61/62/63 editing (fig. 6C, bottom left). The low expression level of proteins containing the US2L signal peptide (fig. 6B) suggests that this signal peptide indeed functions inefficiently, which may result in increased sensitivity to translocation defects.

**Figure 6:**
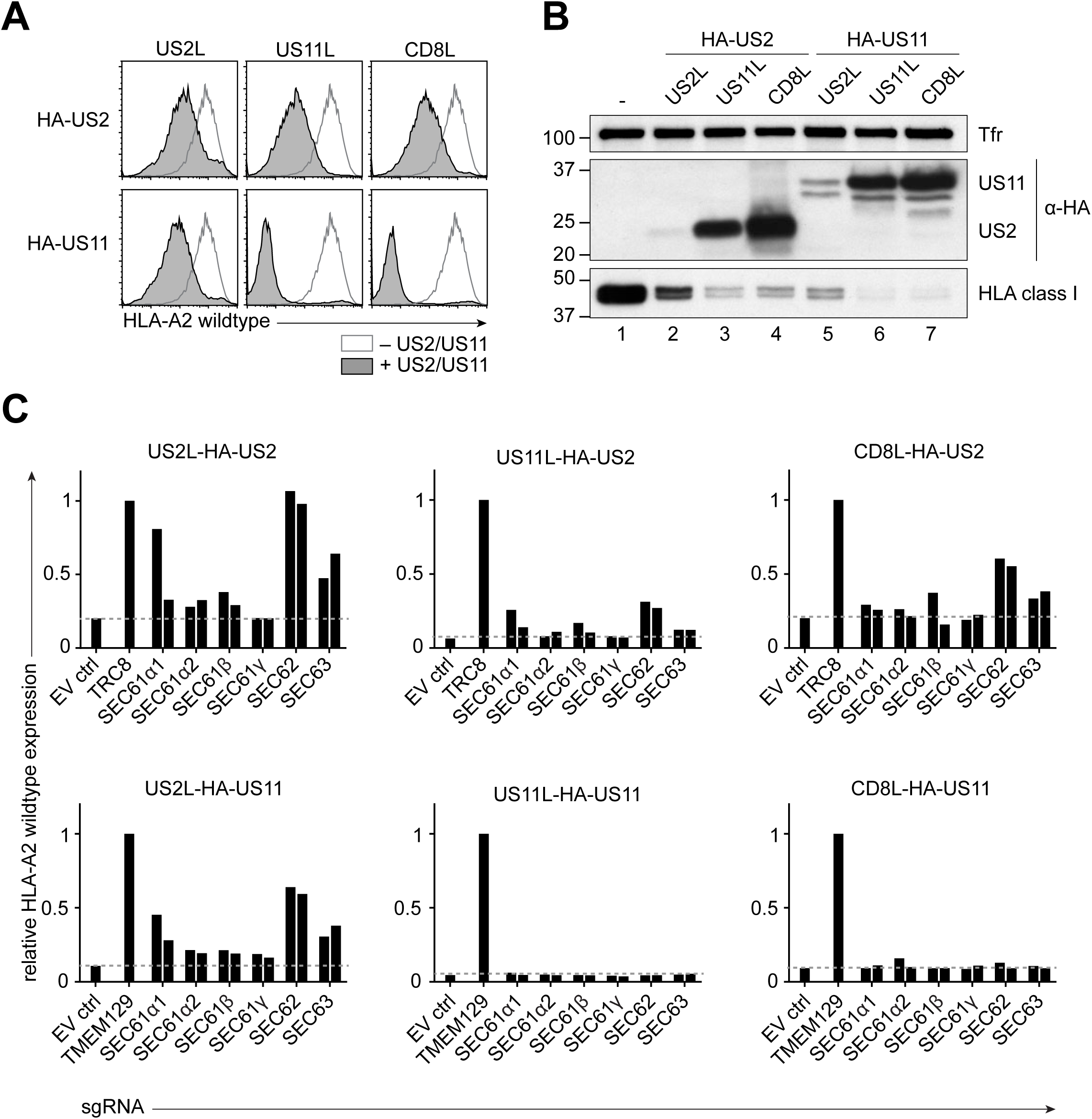
US11 becomes dependent on functional SEC61/62/63 when fused to the US2 signal peptide. A) The US2 variants from figure 5A as well as HA-US11 variants containing the three different signal peptides were expressed in U937 cells containing a non-tagged HLA-A2 construct. All HA-tagged US2- and US11-variants downregulate this HLA-A2 molecule, as shown by flow cytometry on cell surface expression of HLA-A2 using the allele-specific BB7.2 antibody. B) Western blot on the cell lines from A. Protein expression levels of HLA-I, US2/US11 (via their HA-tags) or Transferrin receptor (TfR; loading control) were assessed. HLA-I expression levels reflect the flow cytometry data shown in A. The expression levels of US2 and US11 are affected by their signal peptides, with US2L leading to a low, US11L to an intermediate and CD8L to a high expression level. C) Relative wildtype HLA-A2 expression in the presence of SEC61/62/63 sgRNAs. This experiment was set up similarly to figure 5B, albeit with the untagged HLA-A2 construct. HLA-A2 expression was assessed at the cell surface using an HLA-A2-specific antibody. Flow cytometry data is summarized in bar graphs showing the HLA-A2 expression level (based on mean fluorescence intensity). HLA-A2 fluorescence intensity is shown relative to the positive control (TRC8 for US2, or TMEM129 for US11) to allow comparison between graphs.

Taken together, HLA-I degradation by HCMV US2 or US2L-containing US11 is decreased when subunits of the SEC61/62/63 complex are edited by CRISPR/Cas9. Our data suggest that the inferred dependence of US2 on the SEC61/62/63 complex is specific to the inefficient signal peptide it bears.

## Discussion

In this study, we assessed the role of SEC61, SEC62, and SEC63 components in HCMV US2 translation. To study the role of the SEC61 complex in US2 functionality, we established clonal cell lines containing mutants of single subunits of the SEC61 complex. Clonal cell lines provide a well-characterized context to study the effect of protein mutations or deletions. Although knockdown of certain SEC61 subunits is possible(Lang et al., 2012) and particular mutations in the complex are compatible with life(Haßdenteufel, Klein, Melnyk, & Zimmermann, 2014; Linxweiler et al., 2017), it was surprising that CRISPR/Cas9-mediated knock-outs or severe mutation of these important genes yielded stable clonal cell lines. These cell lines may be used in a broader context for studying translocation.

The characterization of the mutant clones allows us to hypothesize how they affect US2 translocation. For the SEC61α1 clones, the mutations are in the cytosolic loop between transmembrane domains 6 and 7. Crystallography studies suggest that the residues deleted in our cell lines would normally interact with the ribosome(Voorhees, Fernández, Scheres, & Hegde, 2014), which in our cells might lead to a co-translational translocation defect due to disturbed ribosome interaction. Because SEC61α comprises the actual translocation channel and a full translocation block may be lethal to any cell, we hypothesize that the SEC61α1 mutations must have a mild effect on translocation. By contrast, SEC61β is not essential in yeast(Finke et al., 1996; Görlich & Rapoport, 1993). In line with this, we have been able to create clonal full knockouts of SEC61β. However, SEC61β’s non-essential role in translocation suggests that translocation could still occur in our knock-out cells. We also confirmed the previously reported interaction between SEC61β and US2(Wiertz, Tortorella, et al., 1996) (data not shown). In contrast to SEC61β, SEC61γ is essential in yeast(Falcone et al., 2011). The relatively small HLA-I rescue phenotypes observed upon SEC61γ editing resemble those of essential genes like SEC61α1 and p97, suggesting that SEC61γ is indeed important for cell survival. Alternatively, SEC61γ might simply be less important for US2 function. SEC61γ has however been reported to directly interact with US2(Wiertz, Tortorella, et al., 1996), so mutating it may disturb US2 function and therefore explain the HLA-I rescue we observe. SEC62 has previously been knocked down in mammalian cells(Greiner et al., 2011; Lang et al., 2012), suggesting that it may not be an essential gene for cell survival. We now confirm this with a full knockout clone of SEC62. Interestingly, in SEC62-expressing cells an interaction between US2 and SEC62 can be observed (unpublished observation). The 100-150 amino-terminal residues of SEC62 are required for its interaction with SEC63, through a positively charged domain in SEC62(Wittke, Dünnwald, & Johnsson, 2000). As our sgRNAs target this N-terminal cytosolic domain, we speculate that even the small in-frame deletion observed in clone 1 (around Cys82) may disturb the interaction with SEC63. The SEC63 sgRNAs we use target the gene within its J-domain. This domain interacts with the ER chaperone BiP(Misselwitz, Staeck, Matlack, & Rapoport, 1999). The highly conserved HPD-motif within this domain however remains intact, suggesting that BiP interaction may still occur. Besides being expressed at lower levels, the edited SEC63 in these cells may be less active because of the mutations in the J-domain. Our findings are in agreement with siRNA-mediated silencing of SEC62 and SEC63(Lang et al., 2012). This contrasts the situation in yeast, where Sec62 and Sec63 are essential(Noël & Cartwright, 2018).

Our data suggest that a translocation defect of US2 may underlie the rescue of HLA-I in the SEC61/62/63-edited cells, as we observed a downregulation of US2 upon mutating the SEC genes. This translocation defect appears specific to US2, as the transmembrane transferrin receptor and also HLA-I (in US2-lacking cells) are hardly affected. The sensitivity of US2 to mutated SEC61/62/63 might be two-fold: 1) US2 is inefficiently translocated(Gewurz et al., 2002; Wiertz, Tortorella, et al., 1996), making it more sensitive to changes in the translocation process, and 2) as a result of inefficient translocation (even in wildtype cells without SEC61/62/63 editing), US2 is generally expressed at relatively low levels. A further decrease in US2 levels, even when it is minor, may hamper HLA-I degradation when US2 expression falls below a critical threshold. Replacing the signal peptide of US2 by that of US11 or CD8 results in increased US2 expression levels. In cells with highly expressed US2 (or US11) variants, HLA-I degradation is retained even when SEC61/62/63 is edited. This higher protein expression may lead to an excess of US2 or US11, such that minor changes in protein expression due to translocation defects would potentially not affect HLA-I degradation.

The US2 leader is less hydrophobic than most signal peptides(Gewurz et al., 2002), while this hydrophobicity is crucial for its association with SRP. The low affinity of US2 for SRP may strongly influence the efficiency of co-translational translocation. The US2 signal peptide does not contain any rare codons, which may result in the translating ribosome having an insufficient time window to engage SRP, thereby lowering the chances of successful translocation even further. Finally, the US2 signal peptide also contains a positively charged lysine residue in the n-region, which can divert translation towards the post-translational mode(Guo et al., 2018). The low hydrophobicity, lack of sub-optimal codons and positive charge in the n-region of its signal peptide make US2 a potential substrate for post-translational translocation.

As post-translational translocation is mediated by SEC62 and SEC63, it was interesting to observe a strong reduction of US2 expression upon CRISPR/Cas9-mediated editing of these genes. In mammalian cells, post-translational translocation is described mainly for protein substrates smaller than 100 amino acids, with 120-160-residue proteins translocation co-translationally albeit in an inefficient manner, with SEC62-mediated translocation as a backup mechanism(Lakkaraju et al., 2012). Its 199-amino acid size makes US2 an unlikely candidate for SEC62/SEC63-mediated post-translational translocation. It would however be interesting to test whether its sub-optimal signal peptide directs US2 towards post-translational translocation. A similar phenomenon has been described for preprolactin, a 227-residue model protein for SRP-mediated co-translational translocation. When the preprolactin signal peptide was mutated, such that it was translocated less efficiently, it suddenly interacted with SEC62 and SEC63(Conti et al., 2015).

## Acknowledgements

We thank Prof. Tom Rapoport and Prof. Domenico Tortorella for vital discussions about the SEC61 mutant cell lines, and Dr. Nicholas McCaul for fruitful discussions about the US2 signal peptide and help with the SignalP data. We would also like to thank the Core Facility for Flow Cytometry in the UMCU for sorting the clonal mutant cell lines. A.B.C.S. was funded by the Graduate Programme of the Nederlandse Organisatie voor Wetenschappelijk Onderzoek (Netherlands Organization for Scientific Research; NWO) (project number 022.004.018).

## Materials & methods

### Cell culture

Human monocytic U937 cells (ATCC) and human embryonic kidney 293T cells (ATCC) were cultured in RPMI 1640 medium (Gibco), supplemented with 5-10% fetal calf serum (BioWest), 100 U/ml Penicillin/Streptomycin (Gibco) and 2 mM Ultraglutamine-1 (Gibco).

### Lentivirus production and infection

The day before virus production, 65,000 293T cells were seeded per well in 24-wells plates. For virus production, 250 ng lentiviral vector was co-transfected with a mix of third-generation lentivirus packaging vectors (250 ng total) using Mirus LT-1 transfection reagent (Mirus Bio LLC). Three days post-transfection, the supernatant was harvested and stored at −80 °C.

For lentiviral transduction, 100 μl virus-containing supernatant supplemented with 8 μg/ml polybrene (Santa Cruz Biotechnology) was added to 30,000 cells. The cells were spin-infected at 1,000G for 1.5 hours at 33 °C.

### Generation of clonal knockout cell lines

To generate knock-out cell lines, U937 cells with stable expression of HLA-A2-GFP, HCMV US2 and *S. pyogenes* Cas9 (Addgene #52962) were transduced with a lentiviral vector containing a CRISPR/Cas9 guideRNA (gRNA) and a puromycin resistance gene. Target sequences of the respective CRISPR sgRNAs are shown below (table 1). Three days post transduction, cells were subjected to 2 μg/ml puromycin to select successfully transduced cells. Single HLA-A2-GFP^+^ cells were FACS-sorted by an Aria3 cell sorter into 96-wells plates containing RPMI 1640 medium supplemented with 50% FCS and allowed to recover. The knock-out status was confirmed by flow cytometric analysis of HLA-I expression, sequencing of the sgRNA target region (table S1), and addback experiments of the respective cDNAs.

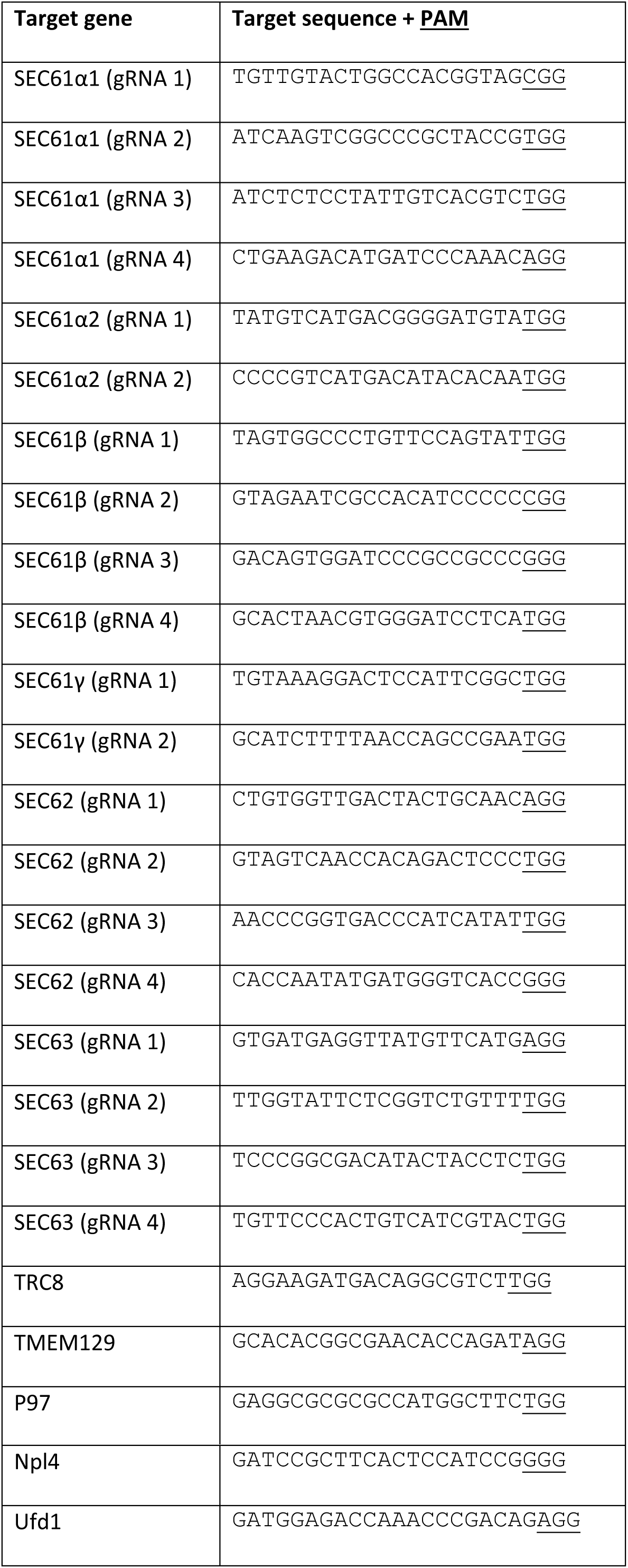

### Antibodies

Primary antibodies used for flow cytometry were: PE-conjugated mouse anti-HLA-A2 (clone BB7.2, BD Pharmingen, no. 558570) and human anti-HLA-A3 mAb (clone OK2F3 LUMC, Leiden, The Netherlands). As secondary antibody for flow cytometry we used PE-conjugated goat anti-human IgG + IgM (H+L) (F(ab’)2) (Jackson Immunoresearch, no. 109-116-127).

For immunoblotting we used the following primary antibodies: mouse anti-HLA-I HC HCA2 mAb, mouse anti-Transferrin receptor mAb (clone H68.4, Invitrogen, no. 13-6800,), rat anti-HA-tag mAb (clone 3F10, Roche, no. 11867423001), rabbit anti-SEC61A mAb (Abcam, ab183046 [EPR14379]), rabbit anti-SEC61B pAb (Abcam, ab15576), rabbit anti-SEC61G pAb (Abcam, ab16843), rabbit anti-SEC62 pAb (Abcam, ab16843), mouse anti-SEC63 pAb (Abcam, ab68550).

Secondary antibodies for immunoblotting were: HRP-conjugated goat anti-mouse IgG (Jackson Immunoresearch, no. 211-032-171), HRP-conjugated goat anti-rat IgG (Jackson Immunoresearch, no. 112-035-175), HRP-conjugated goat anti-rabbit IgG (Jackson Immunoresearch, no. 115-035-174).

### Plasmids

The C-terminally GFP-tagged HLA-A2 construct was expressed from a lentiviral pHRSincPPT-SGW vector (kindly provided by Dr. Paul Lehner and Dr. Louise Boyle, University of Cambridge). Wildtype HCMV US2 was expressed from an EF1A promoter in a lentiviral plasmid. As this plasmid did not contain a resistance gene for antibiotics selection, US2^+^ cells were FACS-sorted as described before (Chapter 4). A lentiviral vector containing an EFS promoter-driven codon-optimized *S. pyogenes* Cas9 fused to a nuclear localization sequence was purchased via AddGene (lenti-Cas9-Blast, plasmid #52962). This plasmid also contains a Blasticidin resistance cassette. CRISPR sgRNAs were cloned downstream of a human U6 promoter in a BsmBI-linearized pSicoR lentiviral vector also containing an EFS (EF1A short) promoter driving expression of a puromycinR-T2A-Cas9Flag cassette. cDNAs were expressed from an EF1A promoter on a bidirectional lentiviral vector. This vector also contains a human PGK promoter driving expression of a Zeocin resistance gene.

### Flow cytometry

Cells were washed in PBS (Gibco) containing 0.5% BSA and 0.02% sodium azide and subsequently fixed 15 min in 0.5% formaldehyde. Cells were stained with the antibodies listed above. Flow cytometry was performed on a FACSCanto II (BD Bioscience) and data was analyzed using FlowJo software.

### Immunoblotting

Cells were lysed in Triton X-100 lysis buffer (1% Triton X-100 in TBS, pH 7.5) containing 10 μM Leupeptin (Roche) and 1 mM Pefabloc SC (Roche). Lysates were spun down at 18,000g for 20 min at 4 °C to remove cellular debris. After addition of Lämmli sample buffer containing DTT, lysates were boiled 5 min at 95 °C and stored at −80 °C until use. SDS-PAGE was performed on 12% polyacryl-amide gels, after which the proteins were transferred to a PVDF membrane (Immobilon-P, Millipore). Primary antibodies were incubated overnight at 4 °C in 1% milk. The secondary antibodies were incubated for 1 hour at room temperature. Each antibody was washed off three times in PBS-T (PBS containing 0.05% Tween20 (Millipore)). Protein bands were detected by incubation with ECL (Thermo Scientific Pierce) and exposed to Amersham Hyperfilm ECL films (GE Healthcare).

### Co-immunoprecipitation

Cells were lysed in Digitonin lysis buffer, containing 1% Digitonin (Calbiochem) in 50 mM Tris-HCl, 5 mM MgCl_2_ and 150 mM NaCl, at a pH of 7.5. This buffer was supplemented with 10 μM Leupeptin (Roche) and 1 mM Pefabloc SC (Roche). Lysates were incubated on ice for 30 min, after which they were spun down at 18,000g for 20 min at 4 °C to remove cellular debris. The supernatant was incubated overnight with StrepTactin beads (GE Healthcare) on a rotary wheel at 4 °C. The samples were washed 4 times with 0.1% Digitonin lysis buffer and eluted with d-Desthiobiotin (2.5 mM d-Desthiobiotin, 100 mM Tris-HCl, 150 mM NaCl and 1 mM EDTA, at a pH of 8.0). 0.45 μM SpinX columns (Corning Costar) were used to collect the eluate. After addition of Lämmli sample buffer with DTT, samples were boiled 5 min at 95 °C and stored at −80 °C until further use. Immunoblotting was performed as described above.

---

**Supplementary table S1:**
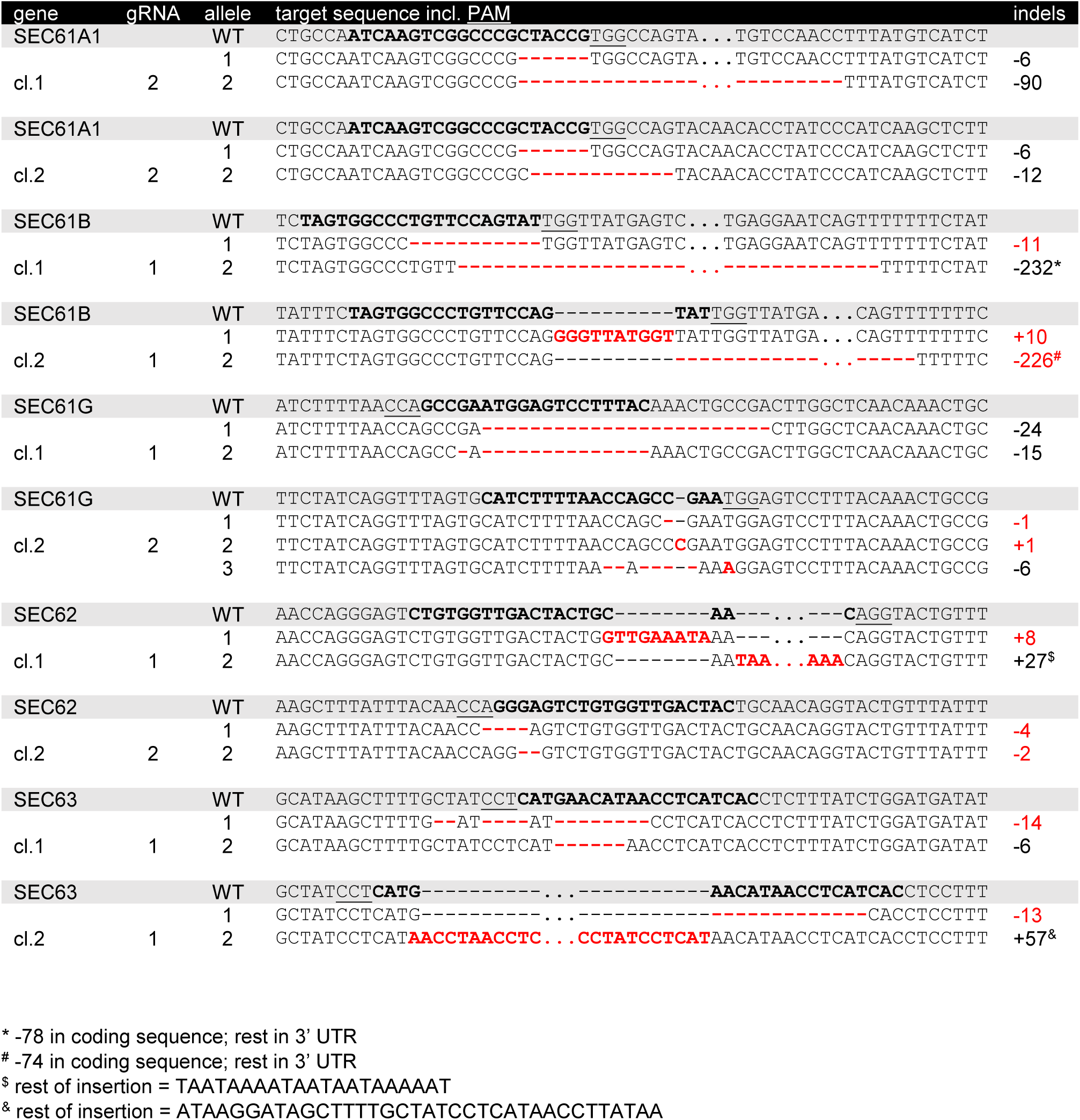
Sequence analysis of clonal mutant SEC61/62/63 cell lines. From all SEC61/62/63 mutant clones, genomic DNA was isolated and the sgRNA-target site was amplified by PCR. Editing of sgRNA target sites was analyzed by standard Sanger DNA sequencing. Only the two selected clones per target gene are shown. SEC61β was edited at the 3’ end of the gene, resulting in a deletion of the C-terminus. A large part of this deletion was in the DNA encoding the 3’ UTR, and did therefore not impact the protein directly. The large insertions observed in SEC62 clone 1 and SEC63 clone 2 are shown only partially. The remaining part of these insertions are listed below the table.

**Supplementary figure 1:**
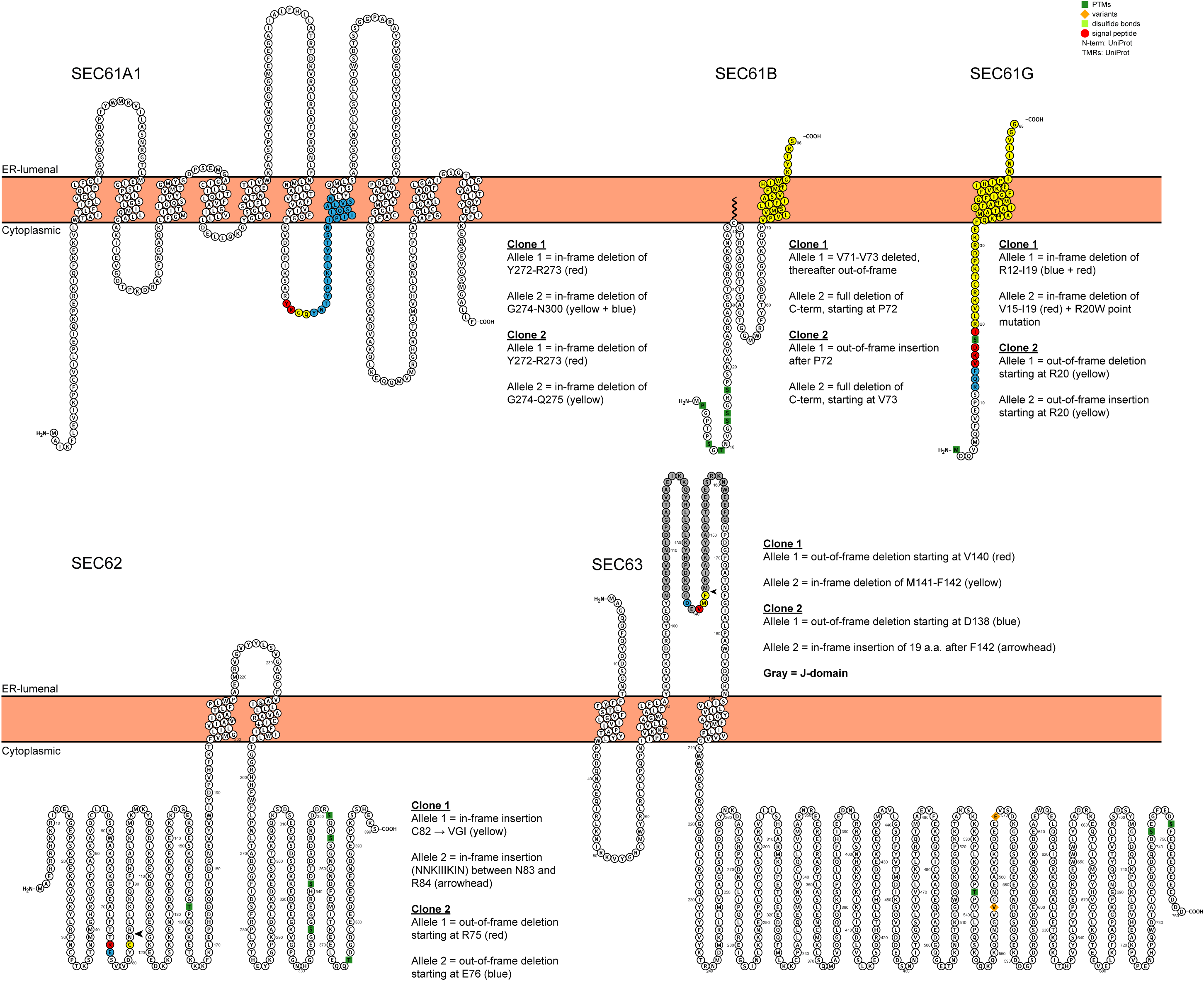
Overview of mutated amino acid residues in clonal SEC-targeted cell lines. Topological view of human SEC61α, -β, and -γ, and SEC62 and −63. Mutated residues are color-coded according to the description in the figure. Topology images are derived from Protter (wlab.ethz.ch/protter).

**Supplementary figure 2:**
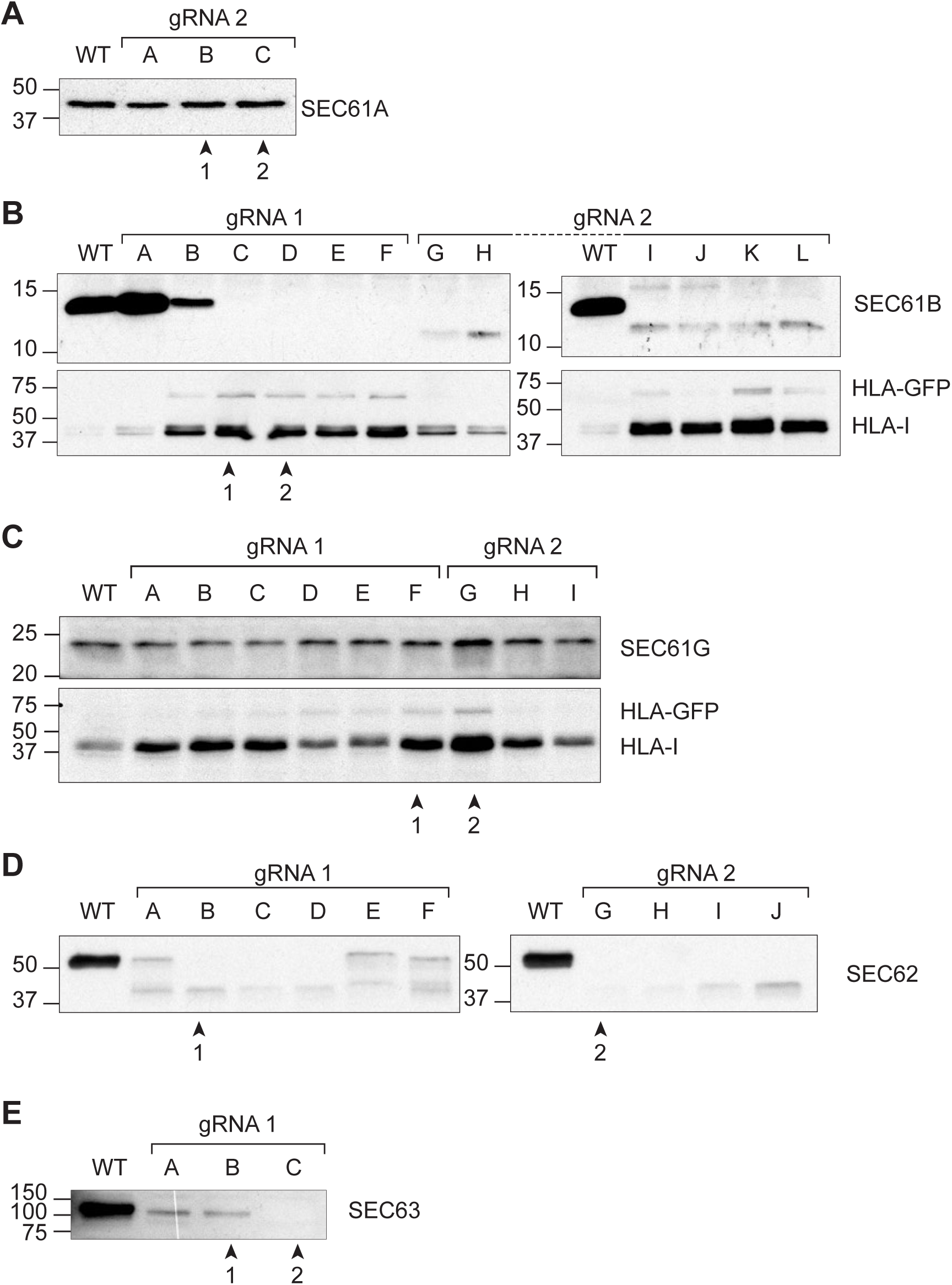
Protein expression of the targeted SEC61/62/63 subunits in clonal mutant cell lines. Immunoblotting was performed for all clonal mutant cell lines generated. Two clones were selected for each target gene, based on non-detectable protein levels in Western blot, the level of HLA-I rescue, or sequencing of the sgRNA target sites in the genomic DNA (table S1). Selected clones are indicated by ‘1’ and ‘2’. Panel A) shows clonal SEC61α1-targeted cells; panel B) those for SEC61β; C) SEC61γ; D) SEC62; E) SEC63. SEC61γ clones were selected based on their HLA-I rescue phenotype.

**Supplementary figure 3:**
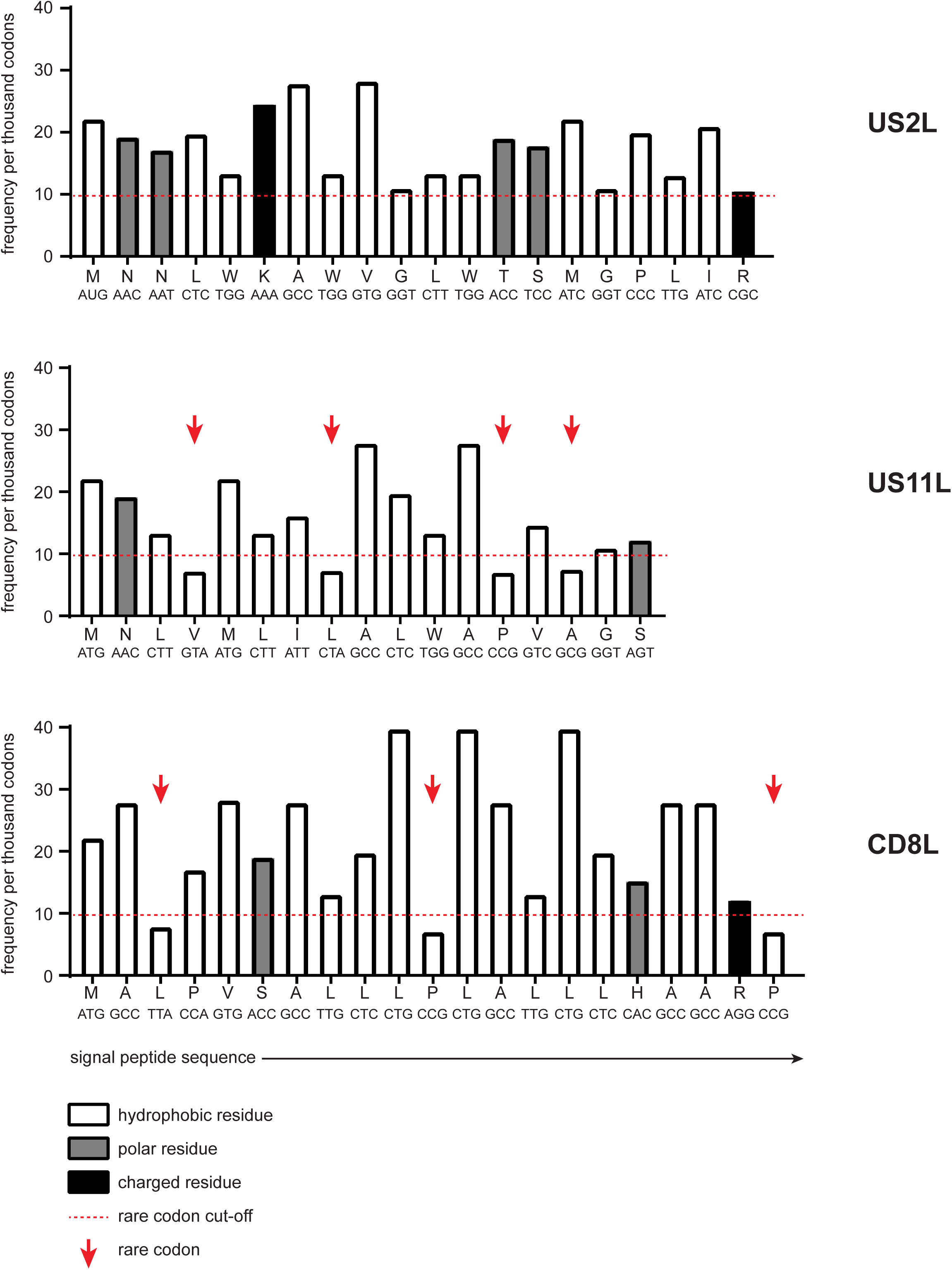
Amino acid and codon properties of the US2-, US11- and CD8 signal peptide. Signal peptide properties of the US2-, US11-, and CD8 leader respectively. On the X-axis, the amino acids and corresponding codons of the signal peptides are shown. The residues are color-coded in the bar graph to show their characteristics with regards to hydrophobicity, polarity or charge. As multiple codons exist for most amino acids, the height of the bars indicates the frequency at which these codons are used (per 1000 codons). These frequencies were derived from https://www.biologicscorp.com/tools/CodonUsage. A cut-off was set at 1% (10 out of 1000 codons), below which the codons are indicated as rare(Kane, 1995).

**Supplementary figure 4:**
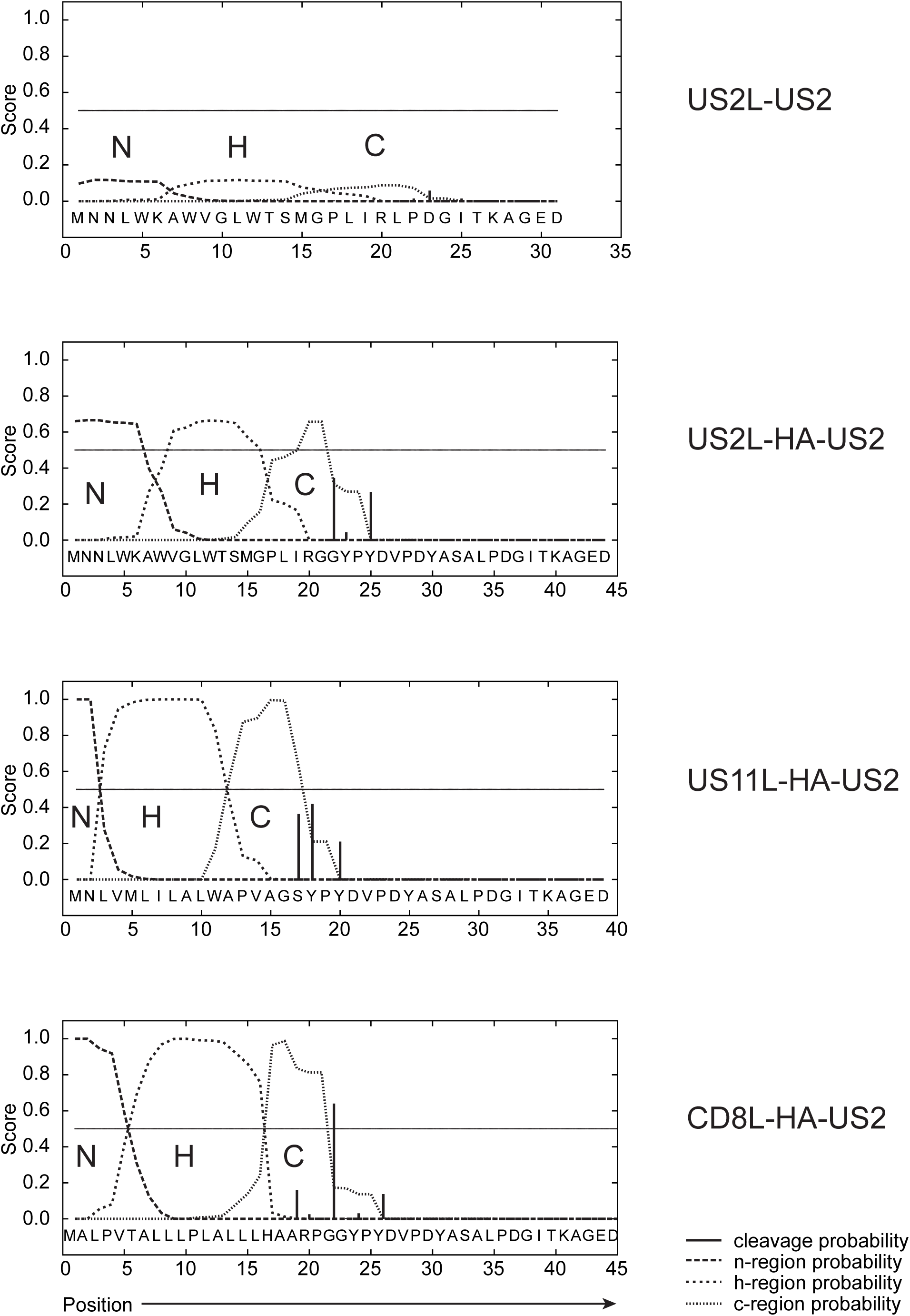
The US2 signal peptide has weak signal peptide characteristics. The N-terminal portions of wildtype US2 or the HA-tagged US2 versions were analyzed using the hidden Markov model (HMM) in SignalP prediction software. This model calculated the probability of the submitted sequence being a signal peptide, including the n-, h-, and c-regions and the predicted cleavage site. For wildtype US2, the signal peptide and part of the protein are shown. For the HA-tagged variants, the signal peptide, HA-tag and the same n-terminal part of the US2 protein are shown. The US2 signal peptide has a low score in this prediction model, suggesting that it bears few of the characteristics of a signal peptide. The HA-tag downstream of the US2 leader may affect signal sequence cleavage, but it is unclear how this impacts the overall HMM score of the signal peptide.

